# Substantial urbanization-driven declines of larval and adult moths in a subtropical environment

**DOI:** 10.1101/2023.10.31.564971

**Authors:** Michael W. Belitz, Asia Sawyer, Lillian Hendrick, Akito Kawahara, Robert P. Guralnick

## Abstract

Recent work has shown the decline of insect abundance, diversity, and biomass, with potential implications for ecosystem services. These declines are especially pronounced in regions with high human activity, and urbanization is emerging as a significant contributing factor. However, the scale of these declines and the traits that determine variation in species-specific responses remain less well understood, especially in subtropical and tropical regions, where insect diversity is high and urban footprints are rapidly expanding. Here, we surveyed moths across an entire year in protected forested sites across an urbanization gradient to test how caterpillar and adult life stages of subtropical moths (Lepidoptera) are impacted by urbanization. Specifically, we assess how urban development affects the total abundance of caterpillars and adult moths, and quantify how richness and phylogenetic diversity of macro-moths are impacted by urban development. Additionally, we determine the effects of urban warming on species-specific adult macro-moth abundance while accounting for urban development and explore how life-history traits condition species’ responses to urban stressors. At the community level, we find that urban development decreases caterpillar biomass and adult moth abundance. We also find sharp declines of adults in response to urban development across the moth phylogeny, leading to a decrease in species richness and phylogenetic diversity in more urban sites. Finally, our study found that smaller macro-moths are less impacted by urban development than larger macro-moths in subtropical environments, perhaps highlighting the tradeoffs of metabolic costs of urban heat island effects favoring smaller moths over the relative benefits of dispersal for larger moths. In summary, our research underscores the far-reaching consequences of urbanization on moths and provides compelling evidence that urban forests alone may not be sufficient to safeguard biodiversity in cities.

## Introduction

Insect declines have been documented across many taxa and regions (reviewed in (Wagner 2020, Wagner et al. 2021), with studies showing declines in insect richness (Forister et al. 2021), abundance (van Klink et al. 2020), and biomass (Hallmann et al. 2017). These declines are particularly alarming given the key role insects play in providing ecosystem services, such as pollination, decomposition, and pest control (Losey and Vaughan 2006, Kawahara et al. 2021). Insect declines are greatest in areas with high human activity (Wagner et al. 2021), and among the many interacting stressors contributing to insect declines, urbanization—a multifaceted form of disturbance—is increasingly recognized as contributing to declines (Fenoglio et al. 2020). The release of local and regional pollutants (air, pesticide, light, and noise pollution), change of ambient temperature due to heat accumulation, and the loss and fragmentation of habitats (Grimm et al. 2008) are all indirect impacts of urbanization. These are all likely to affect insect populations, but the magnitude of declines in urban areas (Egerer et al. 2017, Piano et al. 2020), and consistency of urbanization responses across climatic gradients is still being debated (Secondi et al. 2020).

Urban stressors that influence insect population dynamics also likely interact with species-specific life-history traits, modulating how susceptible populations and species are to urbanization. While some species have shown dramatic declines in the face of urbanization (Merckx and Dyck 2019), others have seen population increases (Raupp et al. 2012). Traits such as body size, mobility, thermophily, and diet generalism are thought to be critical in determining the success of an insect species in an urban environment (Piano et al. 2017, Schmitt and Burghardt 2021). Larger species with greater mobility may allow species to better cope with fragmented urban landscapes (Merckx and Dyck 2019). Species with strong heat tolerance and generalist feeding may also survive under urban stressors in hot cities with low native plant diversity (Merckx and Dyck 2019, Callaghan et al. 2021). However, predicting which life history traits impact urban affinity is challenging, as our knowledge is predominantly based on temperate insect species, which often possess unique characteristics for surviving harsh winters (Wenzel et al. 2020, Theodorou 2022). Therefore, expanding the geographic focus of studies to the subtropics and tropics is critical for better understanding the impact of urbanization on insect populations and community dynamics.

Equally important as expanding geographic foci is extending our understanding of urbanization impacts across insect life-stages. While the effect of urbanization on the abundance and diversity of adult insects has been assessed (Fenoglio et al. 2020, Piano et al. 2020, Vaz et al. 2023), larval life-stages have received much less attention, and we are not aware of studies that have simultaneously collected larval and adult data to examine trends across urbanization gradients. In contrast to the growing evidence documenting overall declines of adult moth abundance in response to urbanization (Merckx and Dyck 2019, Gaona et al. 2021), the few studies focusing on caterpillar abundance or biomass have documented increases (Isaksson and Andersson 2007), decreases (Marciniak et al. 2007, Seress et al. 2018), or no evidence of significant trends (Solonen 2001) in urbanized environments. Understanding how insects respond to urbanization across life stages is crucial to conservation planning of insect populations (Radchuk et al. 2013).

Taken as a whole, determining impacts of urbanization on both larval and adult life-stages in subtropical and tropical regions is of pressing priority since these regions host the greatest insect diversity and are areas where urbanization is predicted to expand quickly (Seto et al. 2012). A working hypothesis is that lower latitude insect communities will be more negatively impacted by urbanization in part because they are more sensitive to increases in temperature (Diamond et al. 2015). Two meta-analyses examining if urbanization impacts insects at greater levels in tropical climate regions found contradictory results, highlighting the need for additional evidence. Fengolio et al. (2020) found that the climate region of cities was unimportant in conditioning the effects of urbanization on arthropod diversity and abundance, whereas Vas et al. (2023) reported that tropical zones exhibit a more pronounced negative impact compared to temperate zones. Insect communities in subtropical regions can comprise a mix of species with core ranges occurring in both tropical and temperate zones (Thang et al. 2020), likely making thermal tolerance traits important in predicting species-specific responses to urbanization in subtropical communities (Belitz et al., in review).

Here, we sampled both larval and adult moths across an urban-to-rural gradient for an entire year to test the effect of urban development and urban warming on total caterpillar biomass and adult abundance in a subtropical environment. For adult macro-moths, we also examine the effect of urbanization on richness and phylogenetic diversity. Finally, we test if responses to urbanization stressors differed depending on life history traits for adult macro-moths. We expected increased levels of urbanization to decrease biomass of both caterpillars and abundance of adult macro- and micro-moths. We also expected adult macro-moth abundance and richness to decrease in response to urbanization and for species that are warmer adapted, larger, and less specialized (i.e., caterpillars feed on a greater variety of host plants) to be less impacted by urban development.

## Methods

### Study sites and sampling

We collected adult moths and caterpillar frass approximately once per week at nine study sites along an urbanization gradient in Alachua County, Florida, USA from March 10, 2019, to Feb. 28, 2020. In total, sampling occurred for 51 distinct weeks over this sampling period. The most urban sites were located in the city of Gainesville, a small municipality in North Central Florida, USA with a population of 141,085 (density of 860/km^2^) as of the 2020 census (U.S. Census Bureau 2020). Our rural sites in eastern Alachua County occur in a matrix of intermittent agriculture, semi-natural landscapes, and small towns (< 2,000 residents).

We selected sites by mapping the proportion of impervious surface using the 2016 National Land Cover Database, which provides land cover information at a 30-m resolution (Homer et al. 2020). We included areas classified as developed open space, low intensity, medium intensity, and high intensity development as “developed areas”. Based on the percentage of land classified as developed surrounding each pixel at a 1-km and 10-km scale, we selected three sites each to represent three distinct urbanization classes: urban, suburban, and rural urbanization (Figure 1). Urban sites had at least 60% of the area within 1-km and at least 50% of the area within 10-km classified as developed. Suburban sites had 10-50% of the surrounding land within 1-km and 25-50% of the land within 10-km classified as developed. Rural sites were defined as those where less than 10% of the area around the site was classified as developed at both the 1- and 10-km spatial scales. All nine sites were located within forested conservation areas managed either by the University of Florida, City of Gainesville, Florida Department of Environmental Protection, or local conservation-focused non-profit organizations. Permitting was secured for each site in consultation with the particular land agency administering each site.

**Figure 1.**
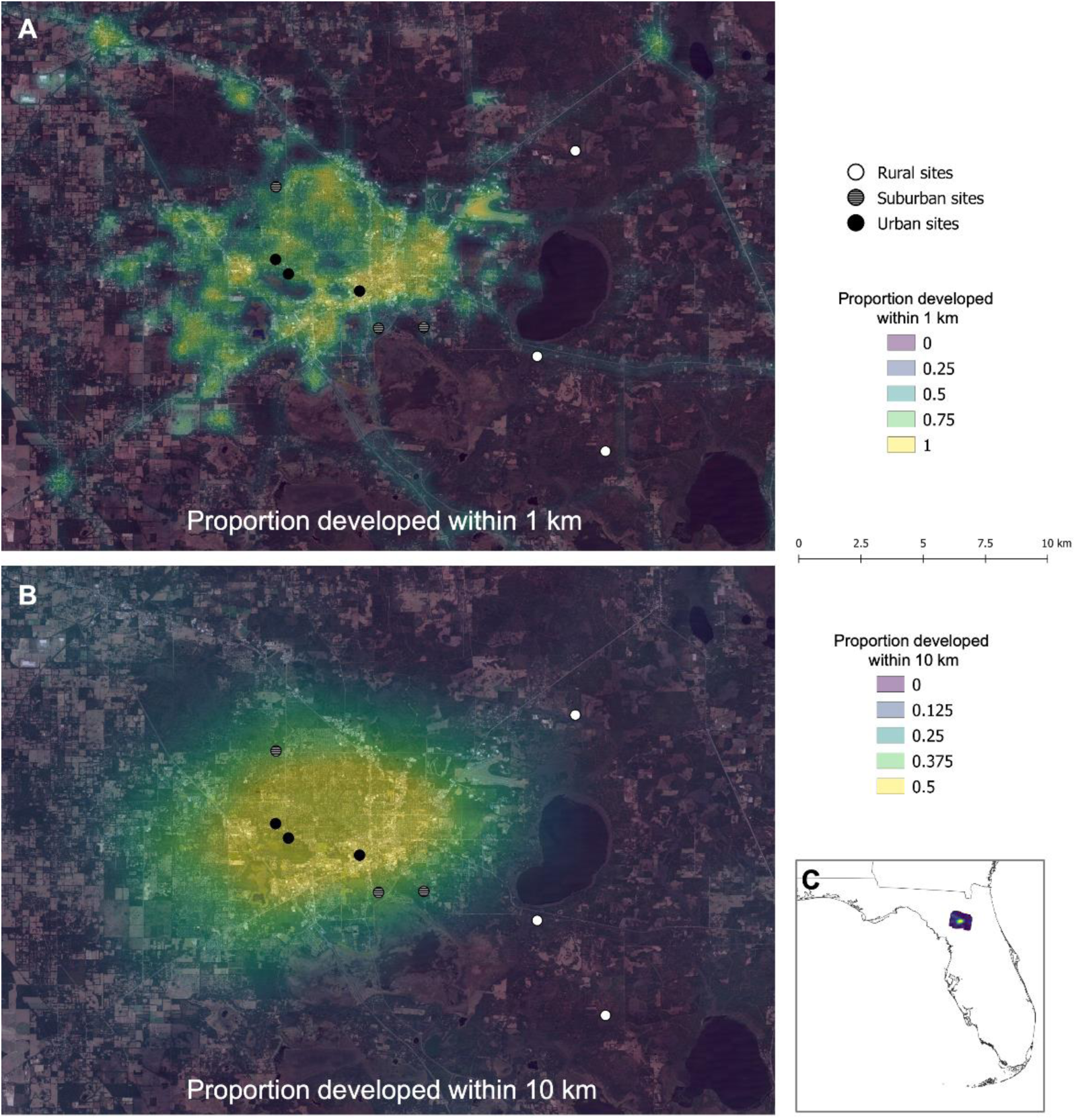
Location of study sites in relation to the urbanization density of Gainesville, FL. The proportion of developed land cover within a 1-km (A) and 10-km (B) buffer is represented by a color gradient. Location of study county within the state of Florida, USA is shown in panel C.

Adult moths were sampled using a single LED funnel light trap per site and caterpillar frass was sampled using six frass traps per site as a proxy for caterpillar biomass (Tinbergen and Dietz 1994). Light traps were built by adapting the low-cost LED funnel trap design described by White et al. (2016) and adding a light sensor to turn on the LED light at dusk and turn off the light at dawn. This trap is known to trap fewer moths than mercury vapor traps but offers a safe, small-battery powered alternative that can facilitate automated trapping in more diverse settings (White et al. 2016). Frass traps were built to sustain sampling throughout the entire wet season. To do so, we built funnels with a radius of 11.66 inches out of wire mesh and attached a plastic collection jar with the bottom of the jar replaced with wire mesh (Figure S1). Mesh funnels were attached to a 20” X 20” wooden frame that was on 24” stilts, allowing the traps to be off the forest floor and above seasonal flooding. We strung a 9” pie tin over the collection jar to serve as a rainfly.

Our sampling protocol consisted of visiting each of the nine sites twice per week. On the first visit, we collected the frass samples from the 6 traps and set the light traps. Any material on the frass trap but not in the collection jar was brushed into the collection jar using a large paintbrush. The collecting jar was then removed and replaced with a new one. Light traps were set by connecting the traps to a 12-volt battery and adding a collecting jar to the trap that was filled with ∼4” of 70% isopropyl alcohol. The following day, the collection jar with insect specimens and the battery were collected. Light trap samples were sorted into groups of lepidopterans and non-lepidopterans, and these samples were stored in 50 mL conical tubes in 70% ethanol (EtOH). Frass samples were dried at room temperature for at least three days in the collecting jars before being sorted, during which non-frass debris, such as plant material and mammal feces, were removed. Once sorted, frass samples were transferred to microcentrifuge tubes where they were stored at room temperature in an HVAC controlled room. After at least 3 months in microcentrifuge tubes, each frass sample was weighed using a scale with 0.001 precision in grams and the amount of frass over the sampling period (mg/day) was calculated.

The total number of micro-moths (defined in this study as moths having a total length of ≤10 mm from head, excluding antennae, to abdominal tip) and macro-moths (total length > 10 mm for the same region) were counted for each light trap sampling day. Additionally, each macro-moth was identified to its lowest taxonomic unit, which was often the species-level.

Species-level identification was not always possible (e.g., due to Geometridae species losing scales, limiting identification to family, or genera like *Datana* where species identification is known to be challenging (Miller et al. 2018). In such cases, specimens were identified to the finest taxonomic resolution possible and were not included in species-specific analyses. We identified all *Halysidota* specimens as *Halysidota tessellaris* even though differentiating between *H. tessellaris* and *harrissii* cannot be done without genitalia dissection. We did so because American Sycamore (*Plantanus occidentalis*), the host plant of *H. harrissii* (Miller et al. 2018), was not observed in the study sites. In total, we collected and sorted 30,497 micro-moths and 5,505 macro-moths.

### Urbanization variables

Two urbanization variables were generated for use as predictor variables in the analysis described below. Those two variables were the proportion of developed land at a 1-km scale (i.e., urban development) and relative temperature of a site (a proxy for urban warming). These variables were weakly positively correlated (r = 0.55). Proportion of developed land reflected the proportion of 30-m resolution pixels classified as developed within a 1-km neighborhood based on the 2016 National Land Cover Database (Homer et al. 2020). To measure the relative temperature of a site, we calculated the temperature of each site relative to the most urban site by keeping one Kestrel DROP D2 (Kestrel Meters, Minneapolis, Minnesota) temperature and humidity data logger at the most urban site throughout our study period and moving a second Kestrel DROP D2 logger to a different site each week. The average difference in mean hourly temperature between each site and the most urban site was calculated.

### Life history traits

We included life history traits as predictor variables in the species-specific adult macro-moth analysis described below. For each identified macro-moth species, we collected the following traits: 1) body size, 2) host plant specificity (HPS), and 3) temperature niche.

Information on voltinism, body size, and host plant specificity were gathered from the *Peterson Field Guide to Moths of Southeastern North America* (Leckie and Beadle 2018). Voltinism (generations per year) was scored as either obligate univoltine or not. Body size measurements were extracted as the upper total length range listed in Leckie and Beadle (2018). In cases where only wingspan was listed instead of total length, body size was calculated as half the upper value of wingspan (García-Barros 2015). Host plant specificity was a categorical variable where species with caterpillars that feed on multiple families or detritus were classified as “generalist”, species that feed on a single family were classified as “intermediate”, and species that feed on a single genus or single species were classified as “specialist” (Futuyma 1976). Temperature niche and temperature niche breadth values were calculated by first downloading occurrence records for each species from the Global Biodiversity Information Facility (GBIF 2023). We then mapped these occurrence records and removed records that fell outside the known range of the species. Using these cleaned occurrence records, we extracted an annual temperature value (using the BIO1 bioclimatic variable available via WorldClim at a 30 second resolution; Fick and Hijmans 2017) for each occurrence point. Temperature niche was the mean value among all annual temperature values.

### Statistical analyses

#### Pooled abundance of caterpillars and adult macro- and micro-moths

We used a hierarchical Bayesian framework using a zero-inflated negative binomial distribution to test the effect of urban development and relative temperature on the total pooled abundance of adult macro-moths and adult micro-moths per sampling event. The non-zero part of the model estimated abundance of adult moths as a function of the proportion of development at a site and the relative temperature of a site. To control for environmental variation in sampling nights, we also included the lunar illumination of the sampling night, total precipitation of the day of sampling, and minimum temperature of a sampling night. Lunar illumination data was gathered using the R package lunar (Lazaridis 2022), and daily weather variables were downloaded from daymet (Thornton et al. 2016). Site was included as a random intercept. The zero-inflated part of the model estimated the probability that a sampling event collected zero moths as a function of the proportion of development at a site, lunar illumination, precipitation, and minimum temperature. Site was again included as a random intercept.

The caterpillar biomass model predicted *log*(*x_i_* + *0.001*), where x = frass mass per site *i* / number of days between sampling events (Seress et al. 2018). Frass mass per day was modeled using a gaussian distribution as a function of the proportion of development at the site, the relative temperature of the site, the average lunar illumination over the collection week, the average minimum temperature over the collection week, and the average precipitation over the collection week. Site was included as a random intercept.

For these models and all models described below we fit models in STAN (Carpenter et al. 2017) using the R package brms (Bürkner 2017) with default priors. In models with multiple predictor variables, continuous predictor variables were scaled to have a mean of zero and a standard deviation of one to allow for easily interpretable model effect sizes across variables. For each model, we ran 2400 iterations each with a warmup of 1000 iterations. No models had divergent transitions (Carpenter et al. 2017) or Rhat values ≥ 1.1. Data simulated from posterior predictive distributions were similar to the observed data. Since our most urban site had by far the lowest abundance in macro-moths, we also ran all models both with the whole dataset and after removing this site to test if our results are robust to the inclusion of this site. Code and associated data to replicate all analyses are archived on Zenodo (https://doi.org/10.5281/zenodo.10056305; Belitz, 2023).

#### Adult macro-moth richness and phylogenetic diversity

For adult macro-moths identified to the species-level, we calculated species richness at each sampling site. We also quantified phylogenetic diversity using Faith’s PD (Faith 1992) and mean pairwise distance (Webb et al. 2008). Species richness was measured as the number of distinct macro-moth species. If a macro-moth was identified to a genus that was not included as a distinct macro-moth species, then those moths were also included as a new “species”. In total, 317 distinct macro-moth species were sampled across our sites.

We calculated community phylogenetic diversity metrics by first generating a synthesis phylogeny for the macro-moth species in our analysis from the Open Tree of Life (Michonneau et al. 2016). Synthesis phylogenies are demonstrated to yield reliable results in community phylogenetic analyses that are similar to purpose-built phylogenies (Li et al. 2019). The database TimeTree of Life (Kumar et al. 2017) was queried to estimate the divergence time of the internal nodes and the branch lengths were scaled from these times using the R package phylocomr (Ooms and Chamberlain 2019). For each site, we calculated proportional phylogenetic diversity as the percentage of overall branch lengths for species found in a site compared to the branch lengths of all species in the total phylogeny (Miller et al. 2018). Abundance-weighted mean pairwise distance was calculated between all species in each site to compare how closely related the average pair of individuals are in a community. Proportional phylogenetic diversity and mean pairwise distance values were calculated using the R package picante (Kembel et al. 2010). We fit a Bayesian univariate linear model using the gaussian distribution to estimate the effect of the proportion of development within 1-km on taxonomic richness, phylogenetic diversity, and mean pairwise distance.

#### Species-specific adult macro-moth abundance

We used a hierarchical Bayesian framework using a zero-inflated negative binomial distribution to quantify the effect of urban development, relative temperature, life history traits, and the interactions among these variables on the abundance of individual moth species at a site. For the positive count data, total abundance of a species collected across the entire year at each site was the response variable.

Predictor variables for the non-zero part of the model were the proportion of developed area at a 1-km scale around the sample site, the relative temperature of each site, moth body size, moth temperature niche, and moth host plant specificity. We also included interaction effects between urban development and body size, relative temperature of the site and temperature niche of a species, and urban development and host plant specificity. A random intercept was included for each species. Additionally, we included a covariance matrix containing the phylogenetic distances between the species as a random intercept term, since ignoring phylogenetic relationships in multi-species models examining trait-environment relationships can lead to overly precise coefficient estimates (Li and Ives 2017). The zero-inflated part of the model estimated the probability that a species was not observed at a site as a function of the proportion of development at a site, host plant specificity, and body size.

## Results

### Pooled abundance of adult macro-moths, adult micro-moths, and caterpillars

At the community level, urban development negatively impacted the pooled abundance of macro-moths and micro-moths, and the biomass of caterpillars (proxied by frass mass) even after accounting for effects of weather and lunar illumination during sampling nights (Figure 2). Adult moths were more abundant in warmer sampling nights, and less abundant in more lunar illuminated nights (Table 1). Caterpillar mass was also higher in warmer weeks, but caterpillar mass was lower in wetter weeks (Table 2). Precipitation did not have a large effect on adult moth abundance (Table 1).

**Figure 2.**
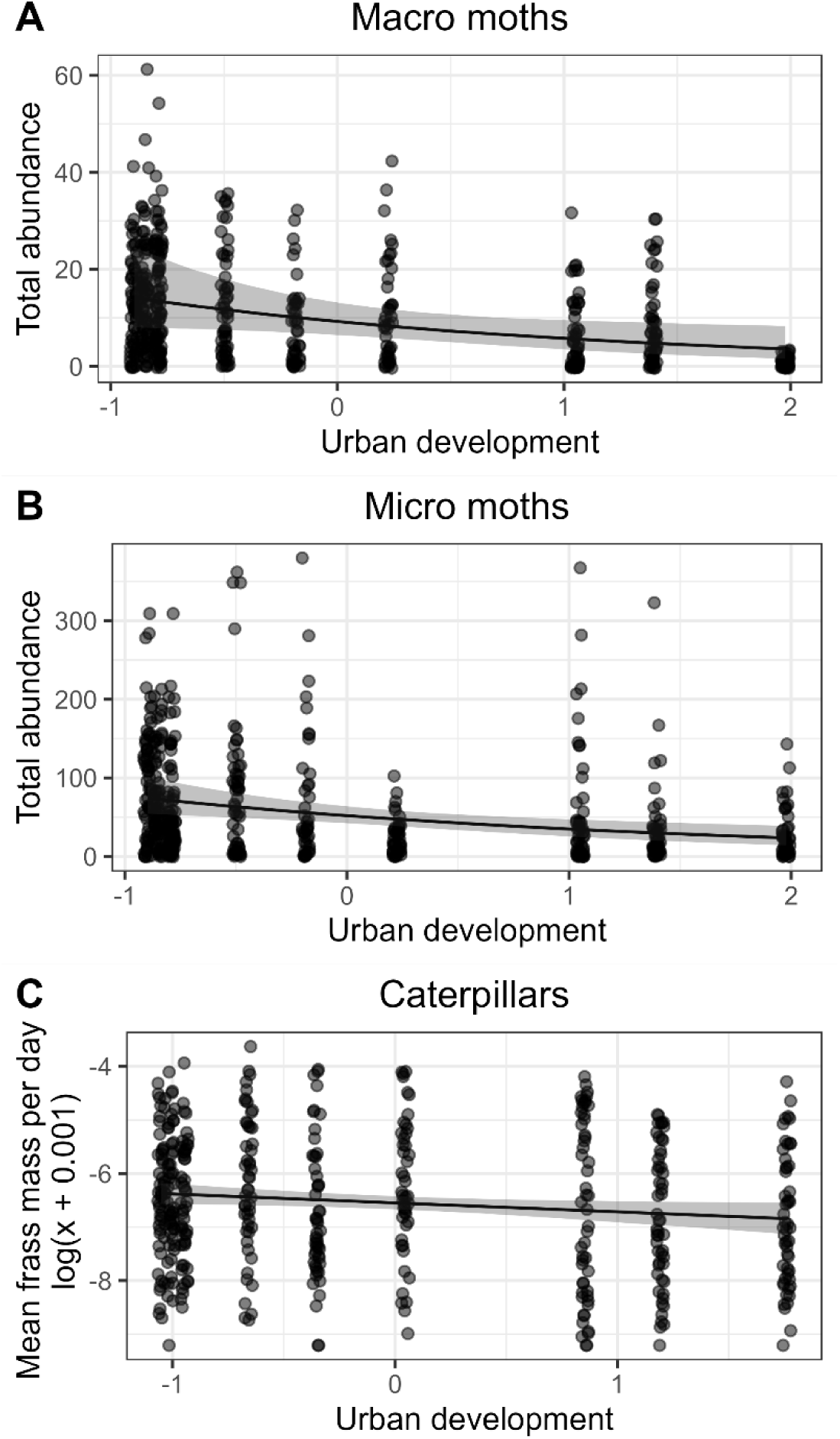
Coefficient estimate and 89% CI of urban development on non-zero abundance of macro-moths, micro-moths, and biomass of caterpillars (as proxied by frass mass). Points represent abundance and biomass collected at individual weeks of sampling for each site.

**Table 1.** Coefficient estimates and 89% credible intervals for the model predicting macro-moth and micro-moth pooled abundance. ZI represents coefficient estimates for the part of the model predicting the probability of zero adult moths sampled.

| Fixed effect | Estimate | <u>Macro Moth</u> |  | Estimate | <u>Micro Moth</u> |  |
| --- | --- | --- | --- | --- | --- | --- |
|  |  | Lower<br>89%CI | Upper<br>89%CI |  | Lower<br>89%CI | Upper<br>89%CI |
| Intercept | 2.23 | 1.86 | 2.58 | 3.95 | 3.75 | 4.16 |
| Urban development | -0.47 | -0.89 | -0.07 | -0.40 | -0.64 | -0.16 |
| Urban warming | -0.26 | -0.68 | 0.15 | 0.16 | -0.08 | 0.41 |
| Lunar illumination | -0.20 | -0.27 | -0.14 | -0.17 | -0.24 | -0.09 |
| Precipitation | 0.04 | -0.05 | 0.12 | -0.04 | -0.12 | 0.04 |
| Temperature | 0.23 | 0.15 | 0.31 | 0.45 | 0.35 | 0.55 |
| ZI Intercept | -6.94 | -10.18 | -4.47 | -9.29 | -12.84 | -6.61 |
| ZI Urban<br>development | 0.77 | -0.09 | 1.67 | 0.10 | -0.70 | 0.86 |
| ZI Lunar illumination | -1.20 | -2.08 | -0.44 | -0.84 | -1.66 | -0.18 |
| ZI Precipitation | 0.35 | -2.60 | 1.30 | -2.80 | -8.33 | 0.21 |
| ZI Temperature | -3.15 | -4.71 | -1.95 | -3.69 | -5.13 | -2.55 |

**Table 2.** Coefficient estimates and 89% credible intervals for the model predicting pooled caterpillar biomass.

| Fixed effect | Estimate | Lower<br>89%CI | Upper<br>89%CI |
| --- | --- | --- | --- |
| Intercept | -6.55 | -6.67 | -6.42 |
| Urban development | -0.17 | -0.32 | -0.02 |
| Urban warming | 0.04 | -0.10 | 0.19 |
| Lunar illumination | -0.05 | -0.14 | 0.04 |
| Temperature | 0.68 | 0.58 | 0.77 |
| Precipitation | -0.24 | -0.34 | -0.14 |

The zero part of our pooled adult abundance models showed that temperature and lunar illumination but not urban development influenced the probability of zero macro-moths being sampled (Table 1). Sampling events were more likely to capture zero macro- and micro-moths on nights that were cooler and had less lunar illumination, but precipitation during the sampling night did not influence the probability of capturing zero adult moths (Table 1). Effect sizes and credible intervals were similar for models predicting total community abundance of macro-moths and micro-moths whether or not the most urban site was included (Table S1). However, in the caterpillar model, the effect size of urban development predicting frass mass was slightly smaller with larger credible intervals (urban development slope coefficient -0.15 [-0.34 - 0.03 89%CI]) in the model that removed the most urban site (Table S2).

### Richness and phylogenetic diversity

Richness and phylogenetic diversity decreased in response to increased levels of urban development (Figure 3). However, the effect size of urban development on phylogenetic diversity is smaller and uncertainty higher when the most urban site is removed from the model (Table S3). We did not find evidence that mean pairwise distance is negatively associated with urban development (urban development slope coefficient estimate -10.29 [89% CI: -25.82 - 5.68]).

**Figure 3.**
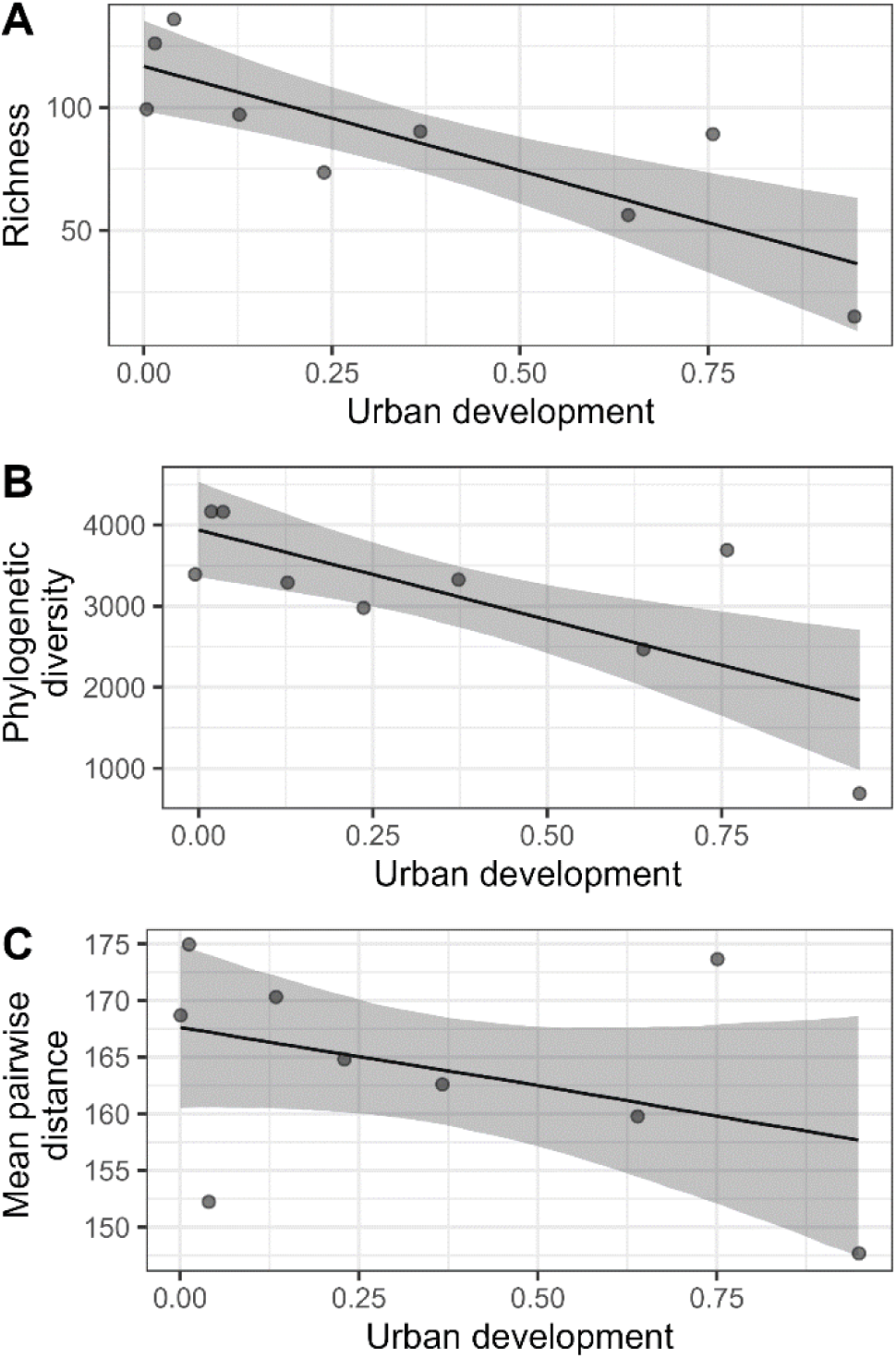
Effect size and 89% CI of urban development on macro-moth species richness (A), phylogenetic diversity (B), and mean pairwise phylogenetic distance (C) .

### Species-specific adult macro-moth abundance

The zero part of our model provides evidence that species were more likely to not be detected (zero abundance) at more urban sites (zero inflated urban development slope coefficient = 5.95 [89% CI: 3.82 - 8.28]). The non-zero part of our model showed that urban development decreases moth abundance, but the body size and host plant specificity of the species mediates this response (Table 3). Specifically, we found larger macro-moths decrease in abundance in response to urban development, while smaller macro-moths had relatively consistent abundance across the urbanization gradient until the most urban sites (Figure 4A). Species that feed on multiple families and that feed on a single genus or species displayed similar decreases in abundance in response to urban development, while species with intermediate host plant specificity showed the greatest urban sensitivity (Figure 4B). Our results also show that relative temperature of a site leads to lower abundance in species with a colder temperature niche (cold-adapted species), but this response has a high degree of uncertainty (Figure 4C; Table 3).

**Figure 4.**
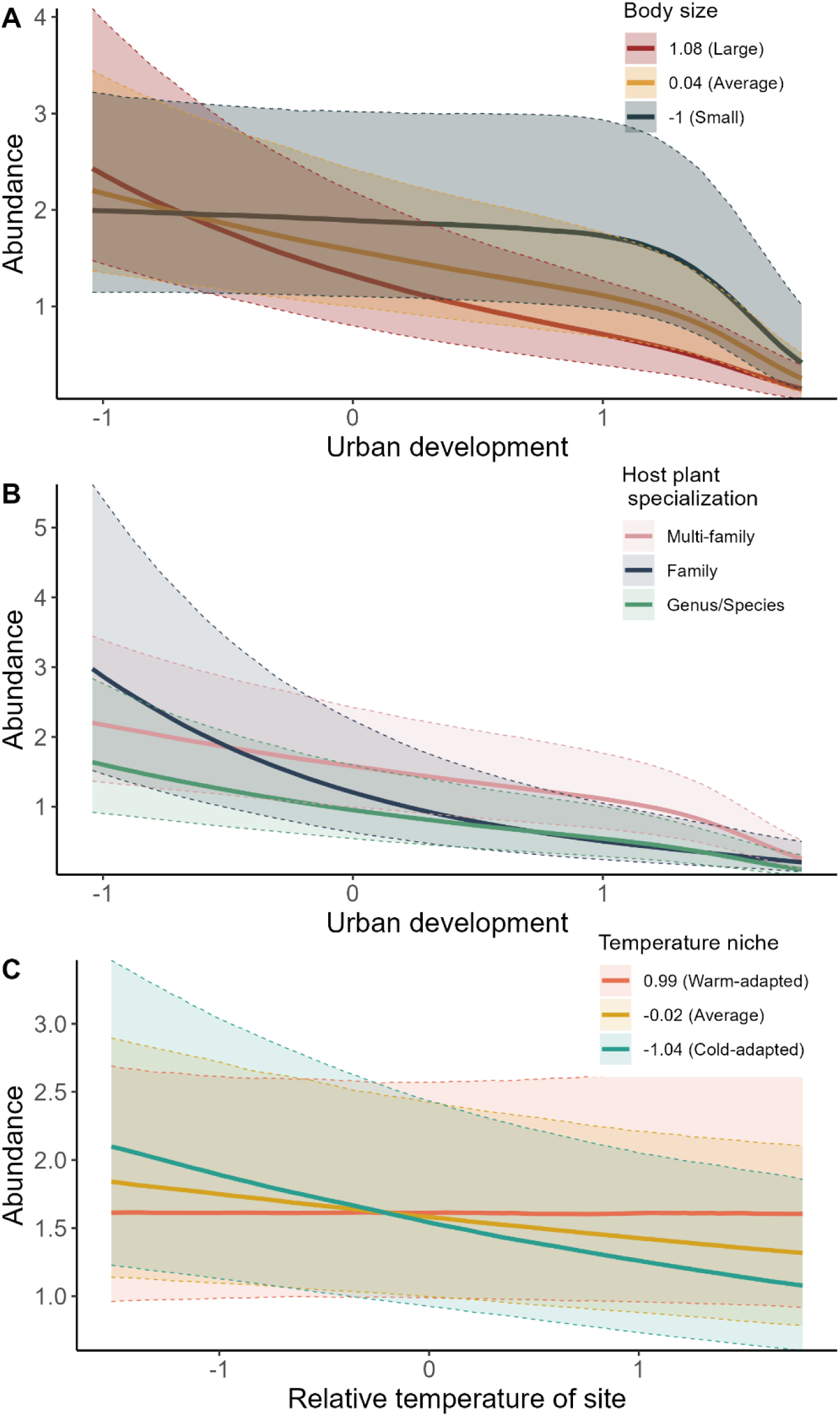
Effect of urban development on species-specific abundance is conditioned by body size (A) and host plant specificity (B). The effect of urban warming is conditioned by a species’ temperature niche.

**Table 3.** Coefficient estimates and 89% credible intervals for the model predicting species- specific macro-moth abundance. ZI represents coefficient estimates for the part of the model predicting the probability of zero adult moths sampled.

| Coefficients | Estimate | Lower<br>89%CI | Upper<br>89%CI |
| --- | --- | --- | --- |
| Intercept | 0.45 | 0.00 | 0.89 |
| Urban development | -0.31 | -0.45 | -0.17 |
| Relative temp of site | -0.10 | -0.21 | 0.01 |
| Body size | -0.17 | -0.40 | 0.06 |
| Temperature niche | 0.02 | -0.16 | 0.20 |
| Host plant specificity (HSP) [2] | -0.27 | -0.79 | 0.27 |
| Host plant specificity (HSP) [3] | -0.50 | -0.97 | -0.03 |
| Urban development:Body size | -0.26 | -0.41 | -0.10 |
| Relative temp of site: Temperature niche | 0.10 | -0.01 | 0.20 |
| Urban development:HSP[2] | -0.55 | -0.88 | -0.22 |
| Urban development:HSP[3] | -0.20 | -0.49 | 0.08 |
| ZI Intercept | -9.81 | -13.66 | -6.47 |
| ZI Urban development | 5.95 | 3.82 | 8.28 |
| ZI HSP [2] | -6.46 | -18.91 | 0.11 |
| ZI HSP [3] | -0.31 | -2.95 | 1.68 |
| ZI Body size | -0.19 | -1.26 | 0.73 |
| sd(Intercept) of phylogenetic relatedness | 0.06 | 0.01 | 0.10 |
| sd(Intercept) of species name | 0.70 | 0.35 | 0.95 |

Phylogenetic relatedness explained little variation in abundance, but the species random intercept was important (Table 3). Overall, the results of the species-specific zero-inflated abundance model remained similar when the most urban site was removed from the dataset, although the effect size of the relative temperature of a site and temperature niche interaction was smaller in the model using the reduced dataset (Table S4).

## Discussion

Urbanization can substantially impact community assemblages by transforming landscapes, including through habitat destruction, fragmentation, increased levels of pollution, and disrupted hydrology (Grimm et al. 2008). In response to urbanization stressors, reductions in both species richness and abundance have been observed across various insect taxa (Fenoglio et al. 2020, Piano et al. 2020), particularly in temperate regions. However, the assessment of urbanization impacts in subtropical regions and their consistency across different life stages and assemblages, such as macro-moths and micro-moths, are far less well known. We conducted repeated sampling of larval and adult moths along an urban-to-rural gradient within a subtropical environment to quantify the effects of urban development and urban warming on larval and adult moth communities. Our study highlights the far-reaching consequences of urban development on moth species and communities, with urbanization negatively impacting the overall abundance of caterpillars, micro-moths, and macro-moths. Furthermore, our study identifies key life history traits that mediate the impact of urbanization on the abundance of individual moth species.

### Adult moth response to urbanization

Adult moths were strongly impacted by urban development with the pooled community abundance of both macro- and micro-moths decreasing along the urbanization gradient. This aligns with the consistent findings in temperate regions, where urbanization has been linked to declines in moth abundance (Bates et al. 2014, Merckx and Dyck 2019, Straka et al. 2021). However, far fewer studies have quantified the consequences of urbanization on insects in subtropical and tropical regions (Wenzel et al. 2020). One of the few studies examining the impacts of urbanization on tropical moths observed that abundance and diversity of moths in the family Geometridae was far lower in urban sites than forest sites (Gaona et al. 2021). The implications of urbanization-driven declines in insect biodiversity in subtropical and tropical environments are particularly disconcerting given the exceptional diversity of arthropods in these regions (Basset et al. 2012) and the projections of expanding urban populations in the subtropics and tropics over the coming decades (United Nations, 2018).

In general, our results demonstrate the prominence of micro-moth diversity in the collected samples, yet identification bottlenecks means that we cannot test phylogenetic and trait driven species-level variation in response to urbanization. This study limitation is not unique to the work presented here, and limits more general predictions of the winners and losers under environmental change. Efficiently identifying micro-moths remains a major bottleneck for ecological studies encompassing entire moth communities. Promising avenues for addressing this issue include automated light traps with computer vision technology and DNA metabarcoding, although the power of these solutions are limited by incomplete DNA and photo libraries (Montgomery et al. 2021).

### The importance of adult life-history traits

Our results indicate that two life history traits are pivotal in identifying the responses of macro-moths at a species-specific level. Notably, larger bodied moths exhibited more negative responses to urban development, while smaller macro-moths showed almost no change in abundances across urban development gradients. These results are contrary to our prediction and a previous study conducted in Belgium that found larger macro-moths were more prevalent in urban sites, which was interpreted as a shift towards increased mobility shaped by habitat fragmentation (Merckx and Dyck 2019). However, our results align with the expectation that the UHIE favors smaller species due to elevated metabolic costs at warmer sites, a pattern observed at the same Belgium-based study sites in non-moth terrestrial arthropods such as ground spiders, ground beetles, weevils, and cladocerans (Merckx et al. 2018). Urbanization may impact biodiversity differently in low-latitude locations, such that in warmer contexts where there is increased heat stress, the metabolic costs of urban heat island effects may favor small moths over the relative benefits of dispersal for larger moths.

Surprisingly, we found species with intermediate host plant specificity were the most impacted by urban development. We expected species with a more specialized larval feeding strategy to fare relatively worse in urban environments, since such a strategy has been identified as an important trait for predicting urban-avoiding Lepidoptera species (Merckx and Dyck 2019, Callaghan et al. 2021) and in animals more generally (Geslin et al. 2016, Callaghan et al. 2019). Urban development can lead to decreases in plant diversity, specifically through reduction of endemic species and increased proportions of exotics (Yan et al. 2019), which will limit opportunities for caterpillars with narrow diets. However, the dominance of oak species, which serve as a host plant for many moth species including specialists (Narango et al. 2020), in all of our forested sites, may have mitigated this effect. Although species with intermediate host plant specificity appear most impacted by the proportion of urban development surrounding a site, we expect specialists to be more broadly affected by urbanization outside of protected forest patches.

A third trait, species’ temperature niche, may interact with urban warming to impact species-specific abundances. Our findings provide some evidence that cold-adapted species experience greater declines in abundance along an urban warming gradient, whereas warm-adapted species maintain stable abundances. This result supports the findings of a previous study that found urbanization to favor thermophilic moth species in temperate regions (Franzén et al. 2020). Our results also extend previous work that found temperature niche to be a key trait predicting which moths will shift their flight timing in subtropical, urbanized environments (Belitz et al. *in review*). It is important to note that the effect size of this result is relatively modest, with sizable credible intervals. Still, taken together, our results, although associated with a degree of uncertainty, suggest that species with colder temperature niches are flying earlier (Belitz et al. *in review*) and have lower abundance in highly urbanized sites.

### Species level losses but lack of phylogenetically clustered filtering

Our results also showcase negative consequences of urban development on macro-moth taxonomic richness and phylogenetic diversity. Our results corroborate recent work indicating that streetlamps with UV emission negatively affect moth species richness on a landscape scale (Straka et al. 2021).

However, we did not find strong evidence of lower mean pairwise distance (MPD) in sites with higher levels of urban development. In sum, these results suggest there is broad filtering of species across all clades rather than clade-specific losses. We might expect MPD to decrease over urbanization gradients if the entire order of Lepidoptera were examined given the fact that the diurnal group of Lepidoptera (i.e., butterflies) may be less impacted by urbanization compared to nocturnal moths (Merckx and Dyck 2019).

### Larval moth response to urbanization

The impact of urbanization on larval insects, such as caterpillars, remains poorly understood, despite their critical roles as herbivores and prey in ecosystems. Our results show a reduction in caterpillar biomass, as indicated by frass fall, across an urban development gradient. Our caterpillar results further strengthen our adult moth findings, since caterpillar sampling avoids the use of light traps which can be susceptible to biases because insects in urban areas may have reduced flight-to-light response (Altermatt and Ebert 2016).

Despite the clear evidence of declines across urbanization gradients, the effect of urban development was less severe for caterpillars than adult moths. One plausible explanation is that moths face high mortality in cities during the transition from late-stage caterpillars to adulthood. Many caterpillars stop feeding near the end of their final instar and wander from their host plant to find a location to pupate (Lee and Roh 2010, Kingsolver et al. 2011). Wandering caterpillars are at greater risk in urban environments, including hazards like road mortality (Ciolan et al. 2017). Moreover, urban soils are often compacted, and topsoil disturbance is common, potentially impeding caterpillars seeking subterranean or leaf litter pupation sites (Schmitt and Burghardt 2021). Those that successfully pupate may still face higher mortality in urban sites due to desiccation, since urban areas are likely to increase dehydration stress (Kaiser et al. 2016) and pupal stages are particularly sensitive to dehydration (Benoit et al. 2023).

We also note that caterpillar and adult abundance proxies, and resolution to taxonomic units, are not the same in this study. Effectively measuring caterpillar abundance or biomass remains challenging as sampling techniques are less developed and field-tested compared to those used for adult moths. Our study takes an important first step at pairing larval and adult datasets, but we recognize that determining the processes underlying differential responses for caterpillars and adults to urbanization are likely complex and continued effort is needed. The ongoing development of environmental metabarcoding techniques offers promising opportunities for using frass traps to determine at least operational taxonomic unit richness and community compositions of forest caterpillars across disturbance gradients (Rytkönen et al. 2019).

### The enormous impact of urbanization on adult moth abundance: conclusions and next steps

Our study reveals a striking pattern where the site surrounded by the most extensive urban development at a 1-km resolution exhibited an order of magnitude lower macro-moth abundance and richness compared to rural sites. In addition to having the most development surrounding the site, this site was also the warmest and had the most artificial light at a 1-km scale (Figure S2).

Nevertheless, the primary conclusions of this study remain robust even when the most urban site was removed from our analysis, reinforcing our finding that urban development has far-reaching ecological consequences on moth communities. The lack of macro-moths at the most urban site suggests the potential existence of ecological thresholds in urban landscapes (Andersen et al. 2009), where abrupt declines in abundance and richness are observed once a certain degree of urbanization occurs. For example, the response of all macro-moths, regardless of body size, decreased in abundance precipitously at the extreme end of our urbanization gradient. Identifying these tipping points and the relative contributions of various urban stressors in reaching them hold crucial implications for urban planning aimed at creating biodiverse cities (Peng et al. 2017). We argue that additional research, including experimental approaches, will be necessary to gain a deeper understanding of the causal mechanisms of urbanization-driven declines in moths (Weisser et al. 2023).

In conclusion, we find extensive consequences of urbanization on nocturnal Lepidopteran communities in a subtropical region, further substantiating that urbanization-induced stressors act at the landscape scale and dramatically alter insect populations and communities. Comparing the rural site with the greatest total abundance and the urban site with the lowest total abundance across the entire year, we documented a 68% reduction in caterpillar frass mass, an 80% reduction in pooled micro-moth abundance, and a staggering 97% reduction in pooled macro-moth abundance. These findings are of particular concern considering that our urban sites were situated within a relatively small city (approximately 150,000 total residents and 860 residents/km^2^) and were located within forested protected parks, highlighting that urban parks alone will not maintain insect biodiversity at comparable numbers to rural areas. Insights from a global meta-analysis suggest that urbanization has a more pronounced impact on insect abundance and richness in tropical areas compared to temperate regions (Vaz et al. 2023). This phenomenon may in part be attributed to the higher baseline abundance and richness found in these areas. Our study raises the possibility that smaller, lower-latitude cities, surrounded by relatively natural areas, may experience larger urbanization-driven reductions in insect biodiversity than larger cities surrounded by more developed landscapes.

## Data availability statement

The data and code to fully reproduce the results and figures presented in this paper can be found on Github (https://github.com/mbelitz/Urban-Moth-Abundance) and is archived on Zenodo (https://doi.org/10.5281/zenodo.10056305). This repository also includes trait information for all species included in our analysis and raw survey data.

## Acknowledgements

This work would not have been possible without a dedicated team of volunteers who helped with processing frass and adult moth samples, and thus we thank K. Shanker, A. Toney, E. Syed, S. Moret, F. Mitchell, N. Federico, C. Kaufmann, and A. Figueroa for their assistance. We also thank CJ Campbell for field assistance and for helping MWB get his car unstuck after a day of field sampling gone awry. Additionally, we thank the land managers, including the University of Florida, City of Gainesville, Florida Department of Environmental Protection, Alachua County Land Trust, and the St. Johns River Water Management District, for allowing us access to their properties for survey activities. I would also like to thank B. Stucky for helping with study design and hardware design for the frass and light traps. Research funding was in part provided by the University of Florida Department of Biology’s Michael May’s Interdisciplinary Grant. MWB was also funded on a fellowship during part of this study by the University of Florida’s Biodiversity Institute.

