## Supplementary material for "Substantial urbanization-driven declines of larval and adult moths in a subtropical environment": Figure S1

### Supplemental Materials

#### Figures

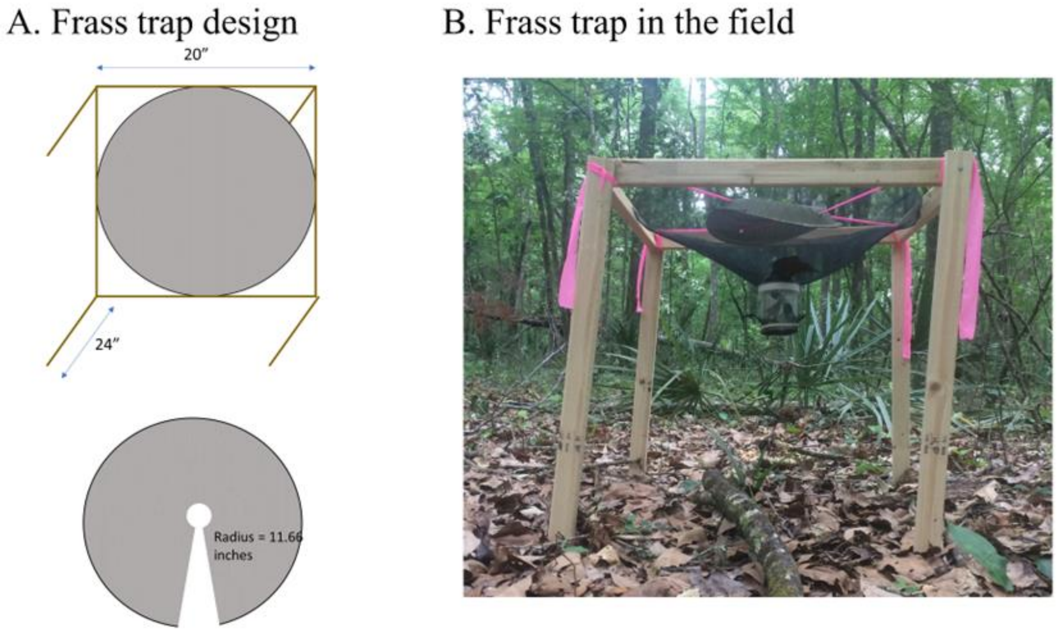


Figure S1. Rain-resistant frass traps were designed with a wire-mesh cone to funnel frass into a well-drained collection jar covered by a rainfly.

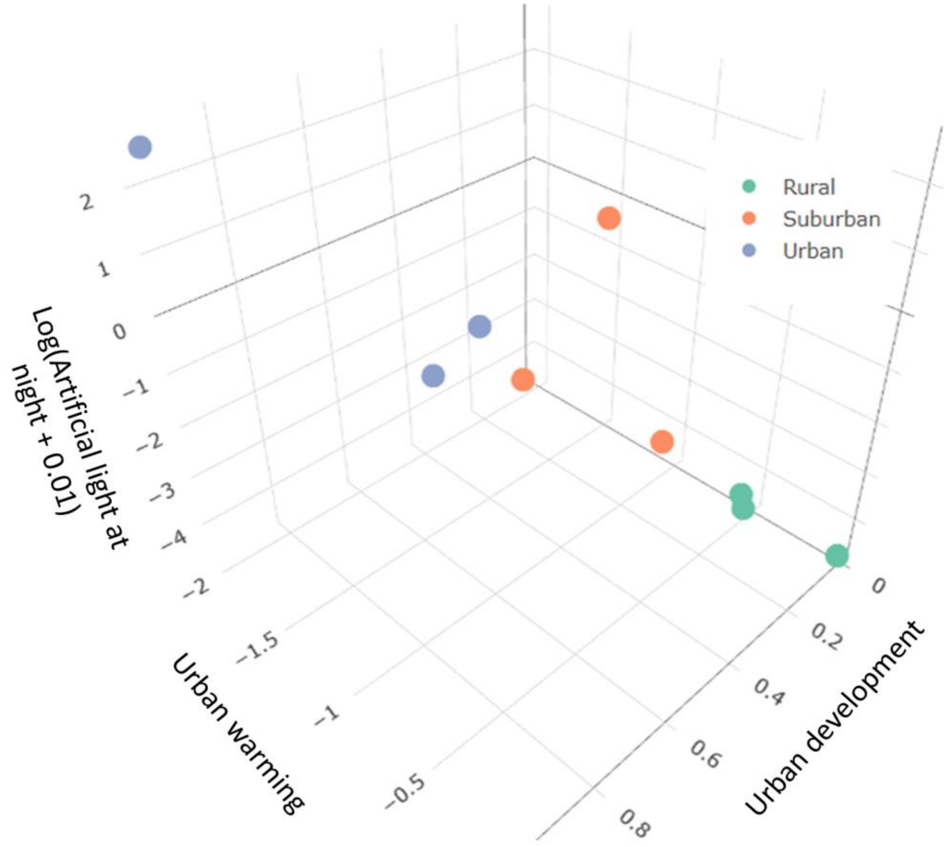


Figure S2. Three dimensional plot showing where sites fall on three axes of urbanization: artificial light at night, urban development, and relative temperature of a site (urban warming). We measured the amount of artificial light around each site by generating 25 random point locations within a 1-km buffer of each site. Light readings were taken using an Extech Light Meter LT300 (Extech Instruments, Waltham, Massachusetts) to the nearest tenth Lux. Light readings were not taken if points occurred on private property or in inaccessible areas.

**Tables**

Table S1. Coefficient estimates and 89% credible intervals for the model predicting macro-moth and micro-moth pooled abundance after removing the most urban site from the dataset. ZI represents coefficient estimates for the part of the model predicting the probability of zero adult moths sampled.

| | Macro Moth | | | Micro Moth | | |
|---|---|---|---|---|---|---|
| Fixed effect | Estimate | Lower 89%CI | Upper 89%CI | Estimate | Lower 89%CI | Upper 89%CI |
| Intercept | 2.38 | 2.25 | 2.50 | 3.97 | 3.72 | 4.22 |
| Urban development | -0.24 | -0.39 | -0.09 | -0.37 | -0.69 | -0.05 |
| Urban warming | -0.02 | -0.17 | 0.13 | 0.21 | -0.10 | 0.52 |
| Lunar illumination | -0.20 | -0.27 | -0.13 | -0.16 | -0.24 | -0.08 |
| Precipitation | 0.05 | -0.04 | 0.14 | -0.03 | -0.11 | 0.06 |
| Temperature | 0.24 | 0.16 | 0.32 | 0.45 | 0.34 | 0.56 |
| ZI Intercept | -6.25 | -9.41 | -4.00 | -9.61 | -13.33 | -6.58 |
| ZI Urban development | 0.73 | -0.29 | 1.77 | -0.11 | -1.21 | 0.91 |
| ZI Lunar illumination | -1.16 | -2.05 | -0.42 | -0.60 | -1.39 | 0.07 |
| ZI Precipitation | 0.22 | -2.85 | 1.20 | -2.78 | -8.46 | 0.35 |
| ZI Temperature | -2.80 | -4.24 | -1.76 | -3.91 | -5.53 | -2.59 |

Table S2. Coefficient estimates and 89% credible intervals for the model predicting pooled caterpillar biomass after removing the most urban site from the dataset.

| Fixed effect | Estimate | Lower 89%CI | Upper 89%CI |
|---|---|---|---|
| Intercept | -6.54 | -6.69 | -6.38 |
| Urban development | -0.15 | -0.34 | 0.03 |
| Urban warming | 0.07 | -0.13 | 0.26 |
| Lunar illumination | -0.06 | -0.15 | 0.03 |
| Temperature | 0.70 | 0.60 | 0.80 |
| Precipitation | -0.24 | -0.34 | -0.13 |

Table S3. Coefficient estimates and 89% credible intervals for the model predicting adult macro-moth diversity metrics. Models fit to data where the most urban site was removed are the models with “No Baca”.

| Model | Coefficient | Estimate | Lower 89%CI | Upper 89%CI |
|---|---|---|---|---|
| Richness | Intercept | 116.80 | 98.67 | 135.32 |
| Richness | Urban development | -84.73 | -124.42 | -45.20 |
| Richness No Baca | Intercept | 111.70 | 94.67 | 128.77 |
| Richness No Baca | Urban development | -59.77 | -104.31 | -14.06 |
| PDP | Intercept | 3938.69 | 3359.57 | 4532.44 |
| PDP | Urban development | -2206.54 | -3532.57 | -952.88 |
| PDP No Baca | Intercept | 3671.76 | 3193.96 | 4130.41 |
| PDP No Baca | Urban development | -927.06 | -2199.37 | 333.85 |
| MPD | Intercept | 167.60 | 160.52 | 174.90 |
| MPD | Urban development | -10.29 | -25.82 | 5.68 |
| MPD No Baca | Intercept | 165.88 | 159.19 | 173.00 |
| MPD No Baca | Urban development | 0.34 | -18.45 | 18.21 |

Table S4. Coefficient estimates and 89% credible intervals for the model predicting species-specific macro-moth abundance after removing the most urban site from the dataset. ZI represents coefficient estimates for the part of the model predicting the probability of zero adult moths sampled.

| Coefficients | Estimate | Lower 89%CI | Upper 89%CI |
|---|---|---|---|
| Intercept | 0.49 | 0.04 | 0.91 |
| Urban development | -0.39 | -0.53 | -0.24 |
| Relative temp of site | -0.09 | -0.20 | 0.01 |
| Body size | -0.19 | -0.41 | 0.03 |
| Temperature niche | -0.01 | -0.18 | 0.17 |
| Host plant specificity (HSP) [2] | -0.27 | -0.80 | 0.28 |
| Host plant specificity (HSP) [3] | -0.50 | -0.97 | -0.03 |
| Urban development:Body size | -0.23 | -0.39 | -0.09 |
| Relative temp of site:Temperature niche | 0.05 | -0.05 | 0.16 |
| Urban development:HSP[2] | -0.44 | -0.80 | -0.09 |
| Urban development:HSP[3] | -0.14 | -0.42 | 0.13 |
| ZI Intercept | -4.60 | -7.69 | -2.64 |
| ZI Urban development | -1.66 | -3.99 | -0.13 |
| ZI HSP [2] | -5.86 | -20.00 | 1.27 |
| ZI HSP [3] | -1.79 | -8.59 | 1.80 |
| ZI Body size | -1.45 | -3.40 | -0.01 |
| sd(Intercept) of phylogenetic relatedness | 0.06 | 0.01 | 0.11 |
| sd(Intercept) of species name | 0.71 | 0.34 | 0.96 |
